# Subcortical source and modulation of the narrowband gamma oscillation in mouse visual cortex

**DOI:** 10.1101/050245

**Authors:** Aman B Saleem, Anthony D Lien, Michael Krumin, Bilal Haider, Miroslav Román Rosón, Asli Ayaz, Kimberly Reinhold, Laura Busse, Matteo Carandini, Kenneth D Harris

## Abstract

Primary visual cortex (V1) exhibits two types of gamma rhythm: broadband activity in the 30–90 Hz range, and a narrowband oscillation seen in mice at frequencies close to 60 Hz. We investigated the sources of the narrowband gamma oscillation, the factors modulating its strength, and its relationship to broadband gamma activity. Narrowband and broadband gamma power were uncorrelated. Increasing visual contrast had opposite effects on the two rhythms: it increased broadband activity, but suppressed the narrowband oscillation. The narrowband oscillation was strongest in layer 4, and was mediated primarily by excitatory currents entrained by the synchronous, rhythmic firing of neurons in the lateral geniculate nucleus (LGN). The power and peak frequency of the narrowband gamma oscillation increased with light intensity. Silencing the cortex optogenetically did not affect narrowband oscillation in either LGN firing or cortical excitatory currents, suggesting that this oscillation reflects unidirectional flow of signals from thalamus to cortex.

**Highlights:** •Local field potential in mouse primary visual cortex exhibits a pronounced narrowband gamma oscillation close to 60 Hz.

•Narrowband gamma is highest in the thalamorecipient layer 4

•Narrowband gamma increases with light intensity and arousal state, and is suppressed by visual contrast.

•Lateral geniculate nucleus neurons fire synchronously at the narrowband gamma frequency, independent of V1 activity.

## Introduction

High-frequency gamma rhythms are produced by a wide range of brain circuits, and are thought to reflect multiple phenomena. Rhythms in a broad gamma range (30–90 Hz) have long been observed in regions including isocortex ^1, 2^, hippocampus ^3^, amygdala, striatum ^4^, and cerebellum ^5^. In cortex, these rhythms are believed to arise from the precisely-timed synchronization of inhibitory networks ^6-9^, and have been implicated in a wide range of functions including coherent transmission of information between neuronal assemblies ^1,10>^, multiplexing of information ^11^, or binding of multiple features of a sensory scene ^12^. Recently, however, it has become clear that gamma rhythms in fact reflect multiple phenomena, and that a single neuronal circuit can support multiple types of gamma rhythm. In hippocampal area CA1, for example, gamma rhythms occurring in distinct frequency bands are coherent with distinct input sources ^13^, perhaps to provide separate routes for information from those structures. Understanding the origin of multiple patterns of high-frequency rhythms is essential for understanding the functional roles these rhythms might play.

In the visual cortex, there is evidence for two different types of gamma-range rhythms, one with power distributed in a broad band between 30 and 90 Hz, and one that oscillates in a much narrower band. The broadband gamma rhythms have long been described in multiple species including cats ^14, 15^, primates ^16-18^, humans ^19^, and mice ^8, 9, 20^. They are modulated by factors such as stimulus position, context, or cognitive state ^14, 17, 18, 21, 22^. A fundamentally different gamma oscillation has been reported in visual cortex of anesthetized cats. This oscillation is extremely narrow in bandwidth in single experiments, and can appear also in the lateral geniculate nucleus and retina ^23, 24^. A gamma oscillation with a sharp band close to 60 Hz has also been observed in visual cortex of awake mice, where its power grows markedly with locomotion ^25, 26^. The origin of this oscillation, and the behavioral and sensory factors modulating it, are unknown.

Here we investigate the narrowband gamma oscillation in awake mouse V1, and establish the factors that determine its strength and frequency, and its relationship to broadband gamma activity. We find that this oscillation has different properties from broadband gamma activity: its amplitude increases with mean light intensity and with locomotion, but it decreases with visual contrast. The narrowband oscillation occurs independently of broadband gamma activity, indicating that it arises from different mechanisms. These mechanisms involve synaptic excitation more than inhibition, and originate before the visual signals reach the cortex. Indeed, the narrowband oscillation is present earlier in the visual system, arising at least as early as the lateral geniculate nucleus.

## Results

We start by characterizing the narrowband gamma oscillation in mouse primary visual cortex (V1). We then explore its synaptic basis and its dependence on visual contrast and locomotion. Finally, we turn to the lateral geniculate nucleus (LGN), and to optogenetic manipulations that investigate the source of the narrowband oscillation.

### Narrowband gamma in V1

Consistent with previous reports ^25, 26^, when we recorded in V1 of awake mice using multi-electrode arrays ^27, 28^ we observed a sharp peak in the local field potential (LFP) power spectrum close to 60 Hz (Figure 1A-C; Supplementary Figure S1). The precise frequency of this oscillation was 61 Hz in this example experiment (Figure 1B-C), and could vary between 55 and 70 Hz in other experiments, with a remarkably narrow bandwidth of 2–5 Hz (Figure 1). This oscillation could not have reflected mains interference, as these recordings were conducted in Europe, where (unlike in the United States) the electricity alternates at 50 Hz.

**Figure. 1.**
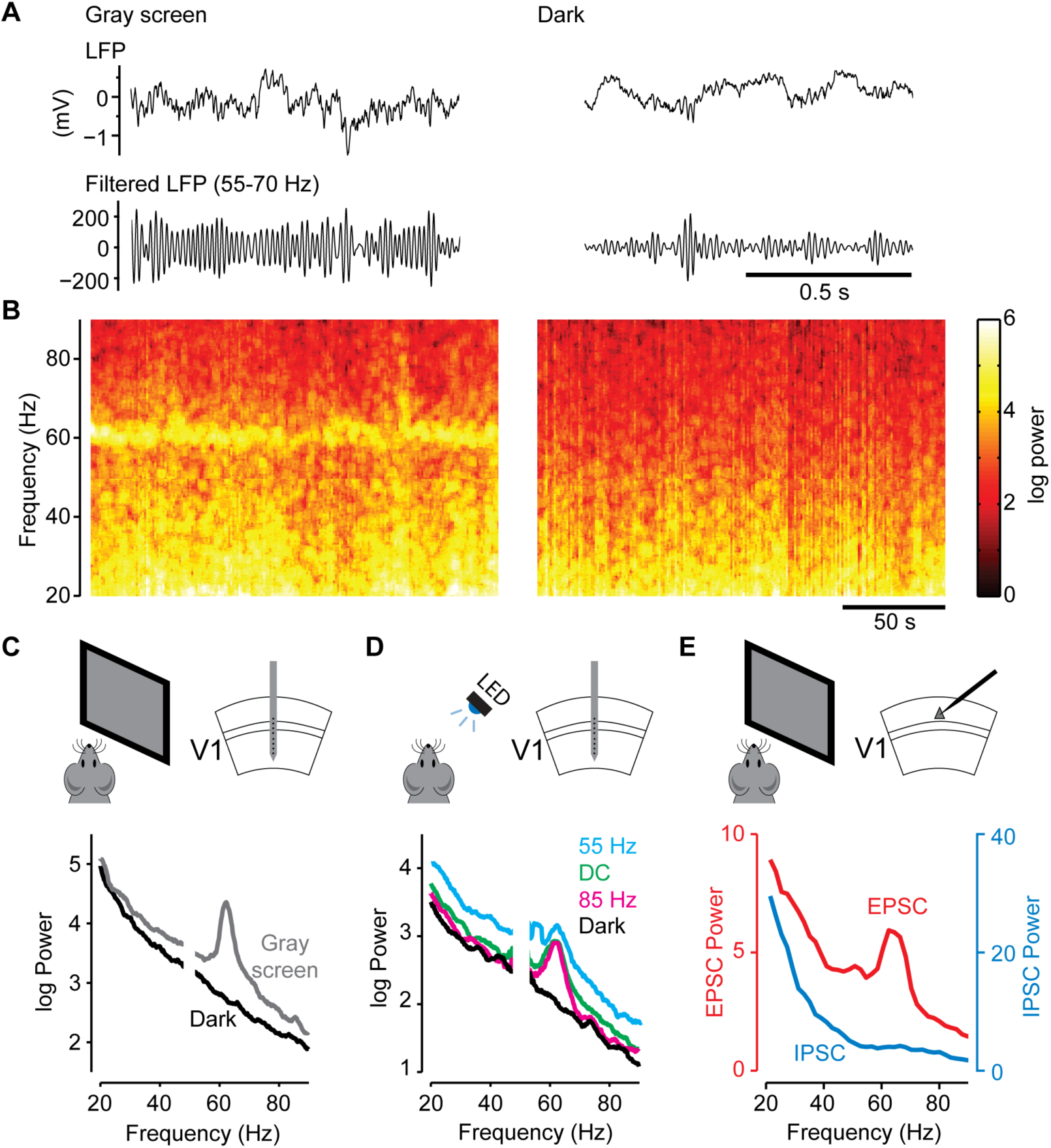
Narrowband gamma in mouse visual cortex. A. Examples of local field potential (LFP) recorded in the primary visual cortex (V1) of mice running on a treadmill while viewing a uniform gray screen (left) or in complete darkness (right). Top traces: 1s of single-trial broadband LFP; lower traces: same data, filtered between 55 and 70Hz. B. Spectrograms showing the variation of local field potential (LFP) power over a longer timescale. The LFP was recorded in the primary visual cortex (V1) of mice running on a treadmill during uniform gray (left) or complete darkness (right). C. Average LFP power spectra during uniform gray screen conditions (gray) or complete darkness (black) (same recording shown in B). D. LFP power spectra under various conditions of LED stimulation: flickering at 55 Hz (cyan), 85 Hz (magenta), constantly on (DC, green), and off (Dark, black) (example from one recording session). E. Power in the excitatory post-synaptic currents (EPSCs, red; whole-cell voltage-clamp recording at −70 mV) and inhibitory post-synaptic currents (IPSCs, blue; whole-cell voltage-clamp recording at +20 mV) in L2/3 neurons in V1, recorded during the uniform gray condition (average spectra across 11 recorded neurons).

This narrowband oscillation was prominent when the mice viewed a uniform gray screen (50 cd/m^2^), but it disappeared when we repeated the experiments in complete darkness (Figure 1A-C; <10^−2^ cd/m^2^). This effect was due to the overall light intensity, not to entrainment of the visual system to the monitor refresh frequency (which can weakly entrain visual cortical neurons ^29, 30^). We confirmed this with control experiments where we stimulated the visual field not with a monitor, but with a light emitting diode (LED, 470 nm) generating light steadily with direct current (DC) input, or flickering at different rates. We saw a narrowband oscillation close to 60 Hz in all cases, independent of visual flicker rates (Figure 1D; p>0.1 t-test for all pairs of flicker rates; see Experimental Procedures).

Unlike broadband gamma rhythms, which involve inhibitory currents ^8, 9, 20, 31^, the narrowband gamma oscillation was primarily mediated by excitatory synaptic currents (Figure 1E). To uncover the synaptic basis of the narrowband oscillation, we analyzed intracellular recordings from layer 2/3 regular-spiking (putative pyramidal) neurons in V1 of awake mice viewing a uniform gray screen^32^ (Figure 1E). Excitatory synaptic currents (EPSCs), isolated by voltage clamping the membrane potential near the reversal potential for inhibition, showed a clear peak around 60 Hz similar to that seen in the extracellular LFP. The power in the narrowband gamma frequency range was significantly greater than a baseline prediction based on a smooth power spectrum that excludes this range (the “residual spectrum”; Supplementary Figure S1; p=0.027; t-test over n=11 recordings). However, we did not observe a similar peak in narrowband gamma power in the inhibitory synaptic currents (IPSCs) recorded from the same neurons during the same stimulus conditions (p=0.11; n = 11 recordings). This suggests that at least in superficial layers, the narrowband gamma oscillation is primarily mediated by excitatory synaptic currents.

### Narrowband gamma is suppressed by visual contrast

Another fundamental difference between narrowband and broadband gamma activity is that increasing visual contrast had opposite effects on them (Figure 2A-D; Supplementary Figure S2). When we measured V1 responses in awake mice to drifting gratings (60^o^ diameter, 2 cycles/sec, 0.05 cycles/°) at different levels of visual contrast, we found a positive correlation of broadband power with contrast *(ρ* = 0.59 ± 0.18, n = 7 recordings; Figure 2A-D). Surprisingly, however, we found that narrowband gamma power was instead negatively correlated with contrast (ρ = −0.43 ± 0.16, n = 7 recordings; Figure 2C,D).

**Figure. 2.**
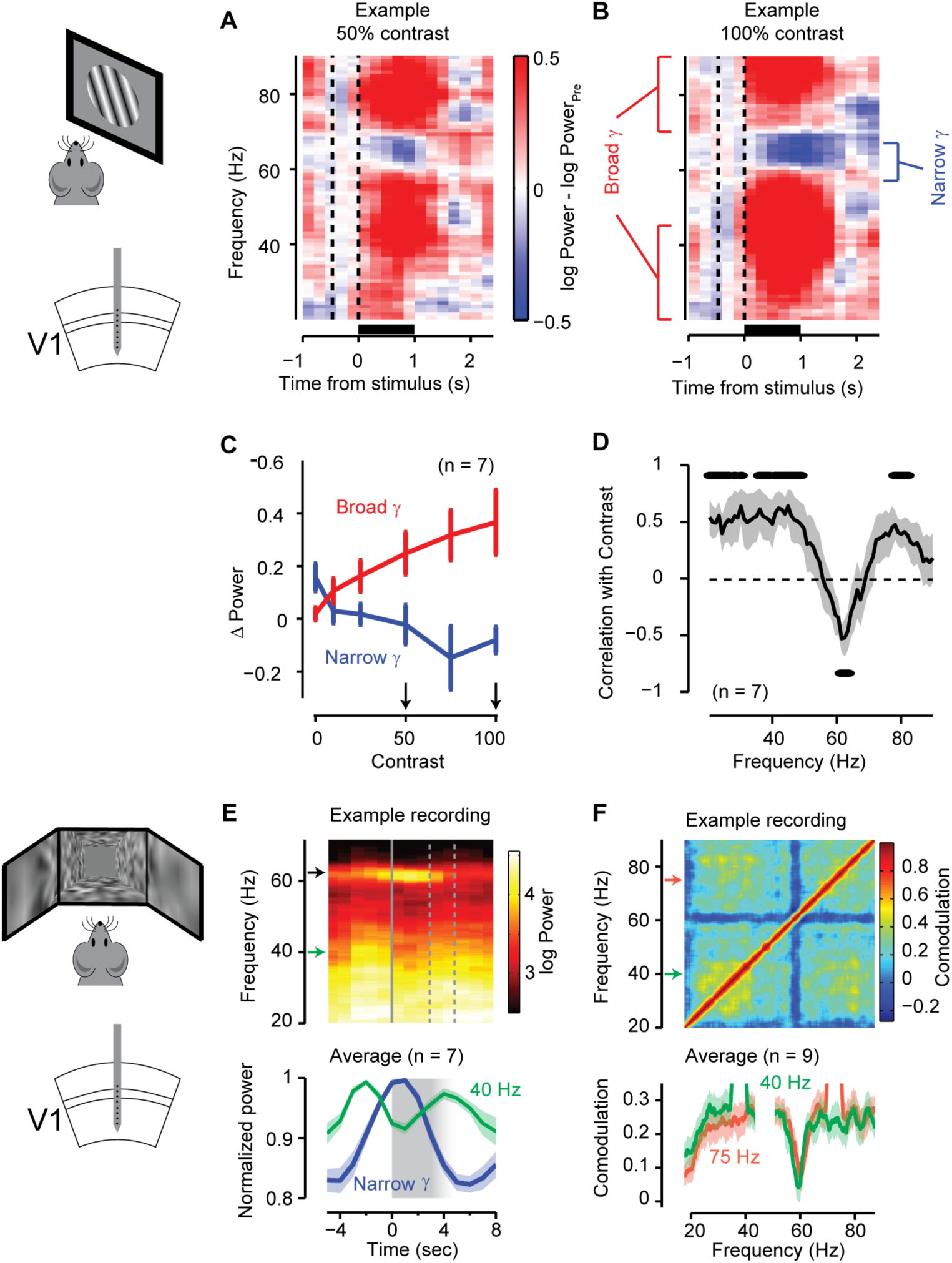
Narrowband gamma oscillations are suppressed by visual contrast. A-B. LFP spectrogram averaged over 20 presentations of a drifting grating stimulus at 50% contrast (**A**) and 100% contrast (**B**), in a single experiment. Color shows power relative to pre-stimulus baseline (-0.5 to 0 s), bars below indicate the period of presentation of a 50^o^ sinusoidal drifting grating in the retinotopic location of the recorded area. **C**. Power at the end of the stimulus presentation period (1 sec) as a function of visual contrast for narrowband gamma (Blue, calculated as the mean across the 55–65 Hz band) and broadband gamma (Red, calculated as the mean across bands 20–40 and 70–90 Hz). The small increase in narrow gamma power for zero contrast (blank) stimuli likely reflects continuing recovery from suppression by the preceding stimulus. **D.** Correlation of power at different frequencies with contrast. Shaded area shows the mean ± s.e.m across experiments (n=7); dots above/below the curve indicate frequencies with significant positive/negative correlations (p<0.05, n = 7). **E.** Average LFP spectrogram triggered on the onset of a gray screen period in a virtual reality environment. The gray screen period lasted between 3–5 secs after which the animal re-enters the high contrast virtual environment. (**Bottom**) normalized power (power / maximal power) at the narrow band frequency (Narrowband gamma; Blue) and at 40 Hz (Green). The shaded regions show the mean ± s.e.m across 7 recording sessions. The gray shaded region shows the period of gray screen presentation, which varied between 2 to 4 s. **F.** Co-modulation of power at different frequencies; i.e. the correlation in the power of different frequency bands, in an example recording. The blue cross visible at 60 Hz indicates that narrowband gamma power is uncorrelated with broadband gamma power. (**Bottom**) Co-modulation of two frequencies of broadband gamma, 40 Hz (green) and 75 Hz (red), with respect to other frequency bands, showing a dip at narrowband gamma frequency. Shaded area shows the mean ± s.e.m across 10 recordings in a virtual reality environment. Arrows in E and F indicate the frequencies shown in the bottom plots.

Narrowband gamma power was suppressed by contrast not only during passive visual presentation, but also while mice used visual inputs to guide locomotion (Figure 2E, F). We recorded V1 activity while mice navigated in a virtual reality environment where visual cues indicated a position that mice must reach to receive a water reward. The narrowband gamma oscillation increased prominently during inter-trial periods, when the screen was uniform gray (Figure 2E; the increase in narrowband power prior to the offset of the previous trial likely reflects the uniform gray texture of the wall at the far of the virtual corridor). Further, throughout the experiment, the power of this narrowband gamma oscillation was uncorrelated with power in broadband gamma range (Figure 2F).

### Narrowband gamma in the lateral geniculate nucleus

The power of the narrowband gamma oscillation varied strongly across the depth of visual cortex, and was markedly higher in layer 4 (L4; Figure 3A,B; Supplementary Figure S3). We recorded the activity across different laminae of V1 using a linear multi-electrode array (16 sites, 50 μm spacing; Figure 3A), and identified layer 4 in each recording as the location showing the earliest current sink^33^ in response to a contrast-reversing checkerboard stimulus (Figure 3A). This layer also showed the strongest ∼60 Hz narrowband gamma peak (Figure 3B). The highest residual power was observed at the same depth as the largest current sink in all recordings (correlation *ρ* = 0.98; p <10^−3^; n = 6 recordings; Supplementary Figure S3).

**Figure. 3.**
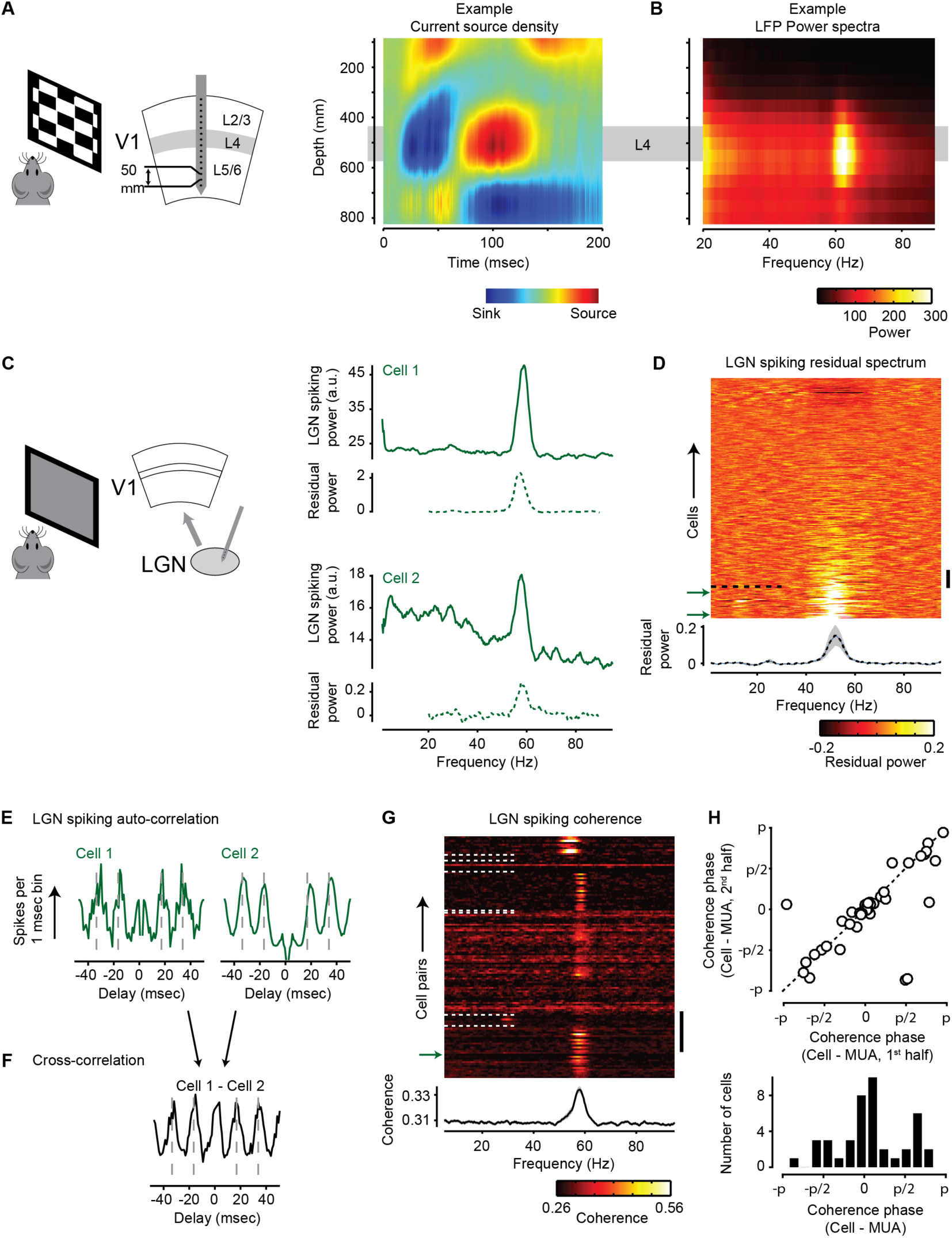
Narrowband gamma is highest in Layer 4 of V1 and present in LGN spiking activity. A. We recorded across the laminae of primary visual cortex using a 16-channel multi-electrode array. Current source density analysis was used to identify layer 4 as the location of the earliest current sink in response to a checkerboard stimulus (figure shows data from one representative experiment). **B.** LFP power spectrum as a function of depth in this same experiment, showing highest narrowband gamma power close to Layer 4. **C.** Extracellular recordings from the LGN using a multi-electrode array in vivo. The spiking power spectra show a clear gamma modulation (two example neurons). Below each spectrum, the dotted line shows the fractional increase in power from a fit to the spectra excluding the narrowband gamma range (residual power spectrum). **D.** Pseudocolor plot showing residual spectra of all recorded LGN neurons (n = 323), ordered by narrowband gamma power. The mean ± s.e.m of the residual power across the population is shown at the bottom. Cells with residual gamma power over 0.20 are below the black dotted line (44 cells). The vertical scale bar indicates 20 cells, and the green arrows point to the examples shown in panels C, E & F. **E.** Two examples of the spike auto-correlation of LGN neurons (same cells as C), showing a clear oscillation at narrowband gamma frequency. Dashed lines indicate the positions of the peaks expected for a 60 Hz modulation (at ±16.7 and ±33.4 msec). F. The cross-correlation of spike times of neurons in G. **G.** Coherence spectra for all pairs of simultaneously recorded neurons with residual gamma power greater than 0.20 (n = 130 pairs). White dotted lines separate pairs from different recording sessions. The mean ± s.e.m of the coherence spectra across all cell pairs is shown at the bottom. The scale bar indicates 20 cell pairs, and the green arrow points to the example cell pair shown in panels F. **H.** (Top) Preferred phase of each oscillatory cell relative to the population of other simultaneously recorded neurons, calculated in the first and second halves of the recording. (Bottom) The distribution of the preferred phase across cells (n = 44 cells).

Because layer 4 receives strong inputs from the lateral geniculate nucleus (LGN), we next asked if the narrowband gamma oscillation was also present in these inputs, and we found that it is evident in many LGN neurons (Figure 3C-D). We used silicon probes to record from 323 LGN neurons from 7 awake head-fixed mice viewing a uniform gray screen (˜50 cd/m^2^). Inspection of spike-train power spectra revealed that many neurons fired rhythmically at frequencies close to 60 Hz (Figure 3C-D). We quantified the rhythmicity of each neuron as the power in the narrowband gamma frequency range, compared to a baseline prediction based on a smooth power spectrum (the “residual spectrum”; Supplementary Figure S1). This measure yielded high values in a sizeable proportion of LGN neurons (Figure 3D). For instance, 14% of LGN neurons (44/323; n = 18 recordings from 7 animals) showed a narrowband gamma peak greater than 20% of the baseline prediction (dotted line in Figure 3D). Consistent with our results in V1 (Figure 2), the power of narrowband gamma in LGN decreased with increasing levels of contrast (ρ = −0.53, p < 10^−4^, n = 9 recordings; Supplementary Figure S4).

The narrowband gamma oscillation entrained the activity of many LGN neurons in a coherent fashion (Figure 3E-H). We observed the narrowband oscillation in the spike time autocorrelations (Figure 3E), and found that it was coherent between simultaneously recorded pairs of neurons (0.33 ± 0.04; mean ± s.d, n = 215 cell pairs where both cells had over 20% residual power), being clearly visible in both the cross-correlogram and coherence spectrum computed from the spike trains of simultaneously recorded pairs (Figure 3F, G). This coherence indicates that the narrowband gamma oscillation jointly entrains the LGN population. Individual LGN neurons had diverse but consistent phases with respect to the oscillation: computing each neuron’s preferred phase in two halves of the data yielded a similar estimate (correlation ρ = 0.69; p < 10^−6^; Figure 3H). The strength and the phase of narrowband gamma entrainment in LGN neurons bore no obvious relationship to their visual preferences, such as receptive field polarity and contrast sensitivity (not shown).

### Narrowband gamma in the LGN is independent of V1

The presence of the narrowband gamma oscillation in LGN suggests but does not prove that the cortex inherits this rhythm from thalamus. Indeed, this oscillation could be generated in V1 independent of LGN activity. To investigate this question, we recorded activity in V1 using extracellular multielectrode arrays, while silencing the activity in the LGN by optogenetically activating thalamic reticular nucleus (TRN) neurons^34^. Silencing LGN strongly suppressed LFP activity in V1 at all frequencies, including the narrowband gamma peak close to 60 Hz (Supplementary Figure S5). This result indicates that the narrowband gamma oscillation cannot be sustained in V1 without input from thalamus.

An second hypothesis is that the oscillation could rely on the presence of cortico-thalamic connections. To test this hypothesis, we asked whether silencing V1 would suppress the narrowband gamma oscillation in LGN, and in the excitatory currents that it elicits in layer 4 cortical neurons. We performed these experiments in VGAT-ChR2 mice, where excitatory V1 activity could be silenced optogenetically across layers^35^ (Supplementary Figure S6). To facilitate combined intracellular and extracellular recordings (Figure 4A), the mice were lightly anesthetized with urethane. This light anesthesia did not impede the narrowband gamma oscillation: LGN neurons showed rhythmic firing at frequencies close to 60 Hz, and EPSCs recorded simultaneously in layer 4 neurons showed a coherent oscillation (Figure 4B, C).

**Figure. 4.**
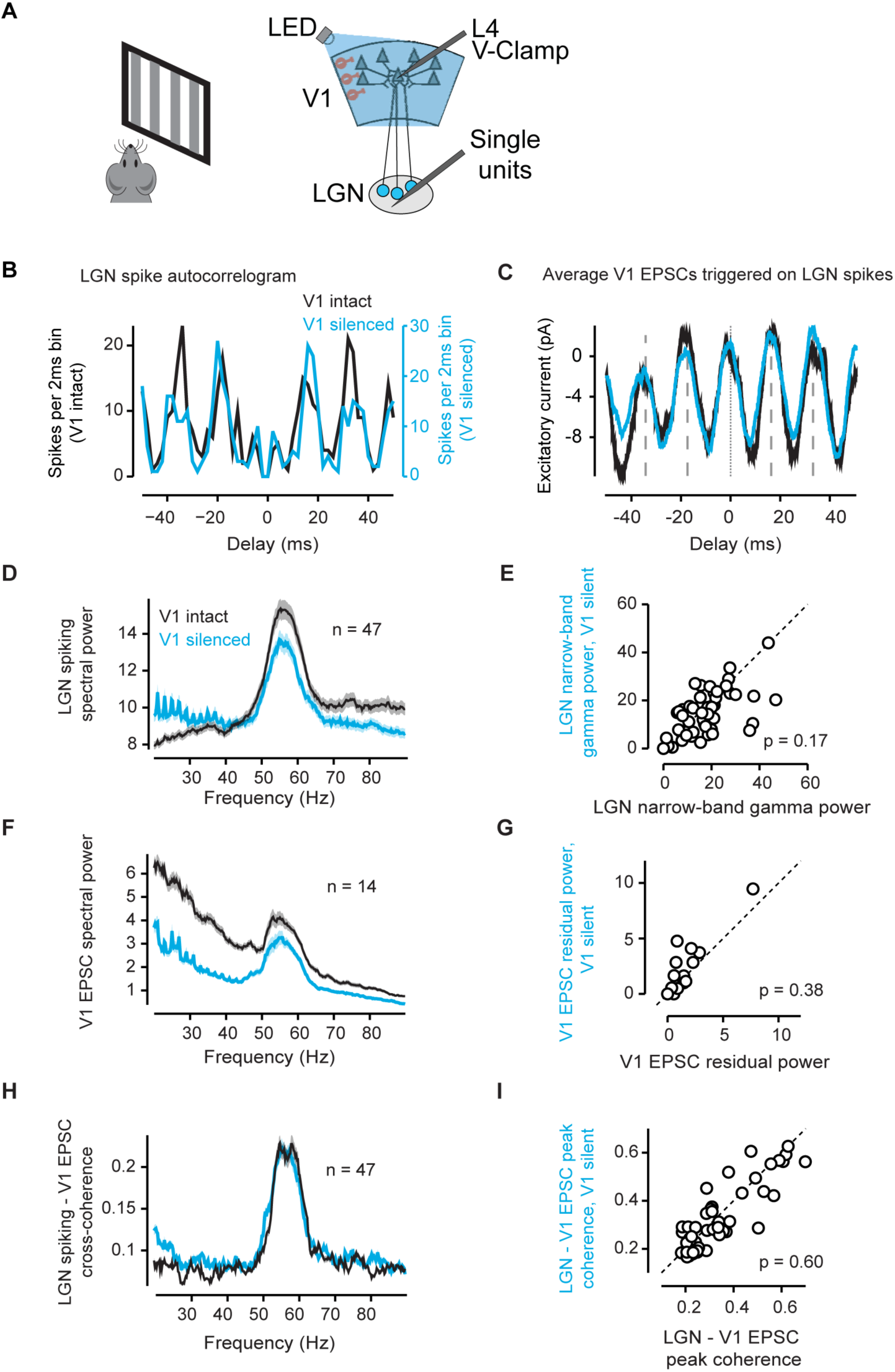
Narrowband gamma oscillations in LGN are independent of V1 activity. A. We made whole-cell voltage-clamp recordings of EPSCs in neurons in layer 4 of V1 while simultaneously recording single-unit activity in the LGN of VGAT-ChR2 mice under light urethane anesthesia. V1 could be silenced by shining a blue LED over the region. **B.** Spike time autocorrelation of an LGN single unit when V1 activity was intact (black) or silenced (cyan) **C.** Average EPSCs recorded by voltage-clamping a V1 layer 4 neuron at −70mV, triggered on spikes of an LGN neuron (same neuron as in B, D), when V1 activity was intact (black) or silenced (cyan). Excitatory current negative. **D.** Mean (± s.e.m) spectral power across LGN neurons (n = 47 neurons). **E.** Comparison of the peak power of the narrow-band gamma when cortical activity is intact or silenced (n = 47 neurons). **F.** Mean (± s.e.m) EPSC spectral power across neurons without and without cortical silencing (n = 14 neurons). **G.** Comparison of residual narrowband gamma power when the cortical activity is intact or silenced (n = 14 neurons). **H.** The cross-coherence between EPSCs recorded in V1 and LGN spiking activity shows a peak at the narrow-band gamma frequency, which does not change with cortical silencing (n = 47 pairs). **I.** Comparison of maximum LGN-V1 cross-coherence when cortical activity is intact or silenced (n = 14 neurons).

Silencing the cortex optogenetically did not affect the narrowband oscillation observed in LGN or the EPSCs in layer 4 neurons (Figure 4). Silencing V1 caused no significant change in the frequency (p = 0.54; paired t-test, n = 47 neurons with greater than 50% residual power in either condition, Figure 4D) or power (p = 0.17, Figure 4E) of the narrowband oscillation seen in LGN neurons. While cortical silencing reduced layer 4 EPSC spectral power at all frequencies, there was no significant change in the residual power of the narrowband gamma oscillation (p = 0.38, n = 14; Figure 4F,G), consistent with addition of a narrowband oscillation inherited from thalamus to cortically-generated broadband activity. The cross-coherence between LGN spiking activity and the EPSCs in layer 4 V1 neurons showed a clear peak at the narrowband gamma frequencies, which was unaffected by cortical silencing (p = 0.60, n = 47; Figure 4H). These results demonstrate that the narrowband gamma oscillation in LGN is independent of feedback from V1.

### Narrowband gamma power and frequency increase with light intensity levels

To investigate how narrowband gamma depends on light intensity, we recorded LGN neurons using extracellular multi-tetrode array recordings while varying the light intensity of a gray screen between ̴1 and ̴95 cd/m^2^ (Figure 5). The light intensity changed every 4 seconds (Figure 5A), presented in interleaved sequences. The narrowband oscillation was seen when light intensities were above 4 cd/m^2^ (Figure 5B,C,E), with power that increased with light intensity (ρ = 0.74, p < 10^−21^, n = 15 recordings; Figure 5B-D). Light intensity also had an effect on the oscillation’s frequency, which increased from close to 50 Hz at low intensities (<5 cd/m^2^) to around 60 Hz at high intensities (>30 cd/m^2^). As a result, we found a strong correlation between the peak frequency of the narrowband gamma oscillation and the log light intensity level (ρ = 0.67, p < 10^−16^, n = 15 recordings; Figure 5E-F).

**Figure. 5.**
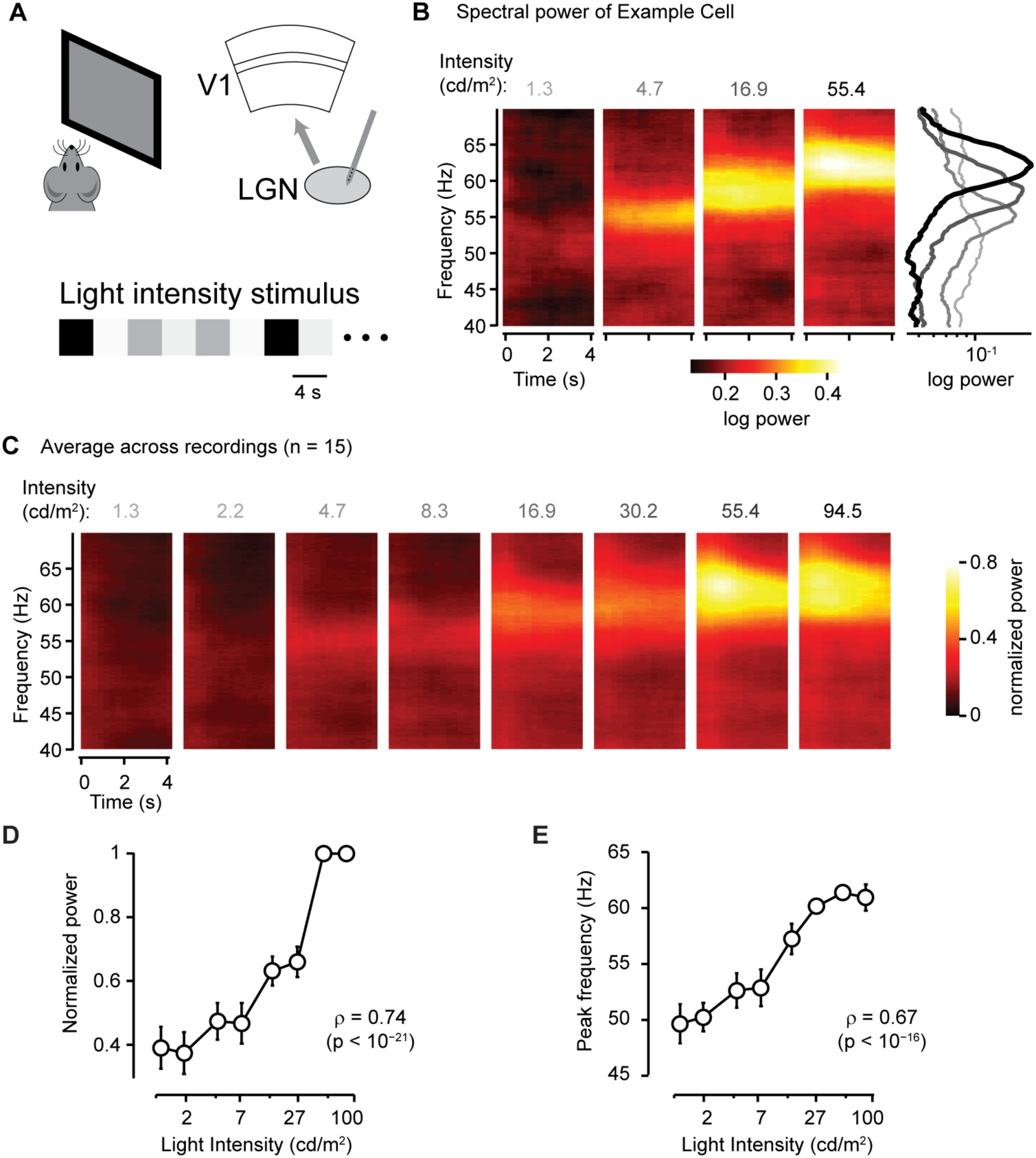
Narrowband gamma power and peak frequency increase with increasing light intensity. A. We recorded activity in the LGN using extracellular electrodes while presenting gray screen stimuli with light intensity changing every 4 s. **B**. Average spectrogram of spiking activity of a single neuron, triggered on the onset of four light intensities (1.3, 4.7, 16.9 & 55.4 cd/m^2^; each intensity was presented 30 times). The mean power spectrum for each intensity is shown on the right (ranging from light gray for 1.3 to black for 55.4). **C**. Average spectrogram of spiking activity across all recording sessions for the 8 different intensities presented, showing an increase in gamma power with intensity (n = 15 recordings). The spectrograms of each recording session were normalized by the maximum average power in each experiment, before averaging across recordings. **D**. Narrowband gamma power is positively correlated with light intensity (ρ = 0.74; p <10^−21^; n = 15 recordings). **E.** The peak narrowband frequency is positively correlated with intensity (ρ = 0.67; p <10^−16^; n = 15 recordings).

### Narrowband gamma is increased in active animals

The strength of narrowband gamma oscillations was related not only to visual factors (light intensity and contrast) but also to the behavioral state of the animal. Consistent with previous reports ^25, 26^, we found that narrowband gamma power is highly correlated with running speed (correlation ρ = 0.49 ± 0.04, mean ± s.e.m, n = 10 recordings in V1). This observation, however, does not imply that behavioral state itself directly modulates gamma power. Indeed, since the mouse pupil dilates during locomotion ^36-40^, and since narrowband gamma power depends on light intensity (Figure 5), it remains possible that the correlation between locomotion and narrowband gamma reflects a greater amount of light striking the retina when pupils are dilated during running.

To answer this question, we measured gamma power after applying the anti-muscarinic agent Tropicamide to the eye surface. This procedure caused full dilation of the pupil (Figure 6A), thus removing the correlation of pupil diameter with running speed. We found that the correlation of narrowband gamma power with running speed persisted even in these conditions of fixed pupil diameter (Correlation *ρ* = 0.35, p<0.01, Figure 6B), confirming that the correlation of narrowband gamma oscillation with locomotion does not simply reflect an indirect effect of increased light flux after pupil dilation.

**Figure. 6.**
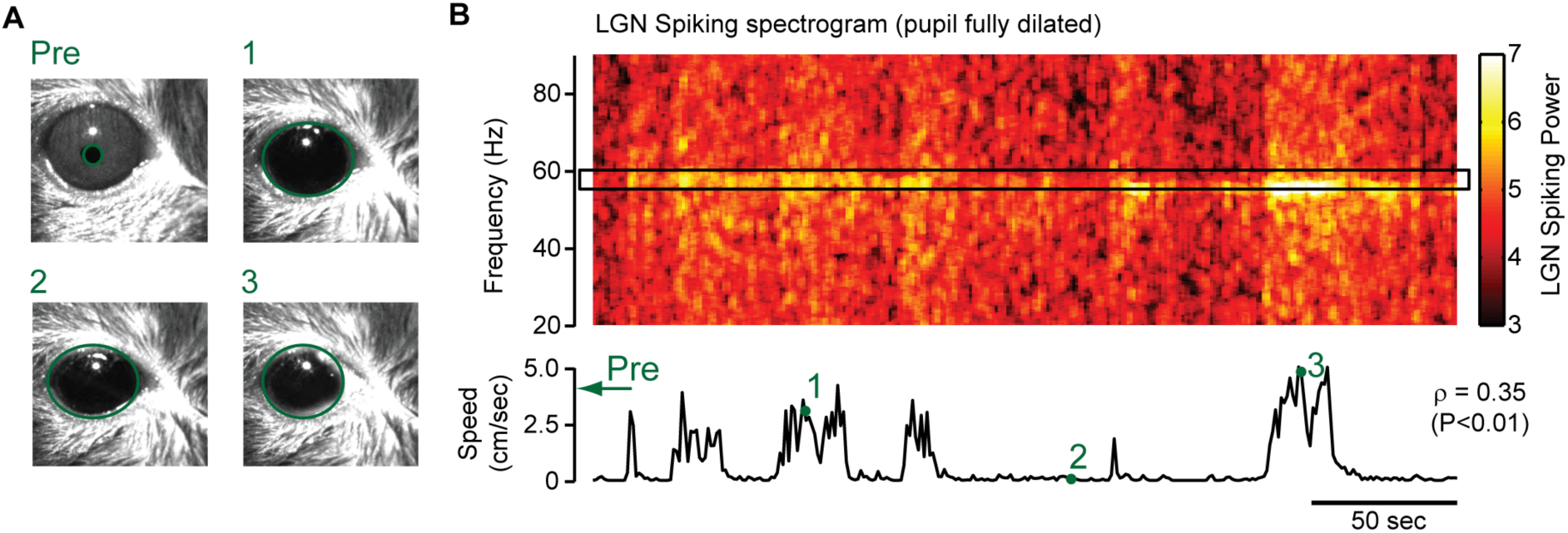
Narrowband gamma power is correlated with running even with a fully dilated pupil. A. Images of the eye before dilation (Pre), and after being dilated (Image 1–3) with Tropicamide (>10 mins after full dilation), when the animal was stationary (Image 2) or running (Images 1 and 3). The green ovals over the images indicate the detected pupil area. **B.** Spectrogram of LGN spiking activity at different frequencies as a function of time. The trace below shows the running speed of the animal. Narrowband gamma power (mean in region highlighted by box) was positively correlated with running speed (ρ = 0.35, p<0.01). The time points of the images are shown in **A** are marked with green numerals.

## Discussion

The narrowband gamma oscillation in mouse V1 has several major differences to the more commonly described broadband gamma rhythm. The power of narrow and broadband gamma activity were uncorrelated, and visual contrast suppressed narrowband gamma oscillation while it increased broadband gamma rhythms. Within cortex, the narrowband gamma oscillation was strongest in L4, and cortical intracellular recordings revealed narrowband gamma oscillation in excitatory, but not inhibitory synaptic currents. Narrowband gamma oscillations persisted in LGN ensembles and L4 synaptic inputs, even after silencing of cortex.

We therefore conclude that the narrowband gamma oscillation represents an entirely different phenomenon than broadband gamma: it is generated in thalamus and possibly even in retina. Whereas broadband gamma rhythms involve cortical inhibitory networks ^6-9^, narrowband gamma is transmitted to cortex by the rhythmic firing of thalamic ensembles. Although resonance in thalamocortical loops has been proposed as a mechanism for the generation of high-frequency oscillations ^41, 42^, we found that the narrowband gamma oscillation was transmitted from LGN to visual cortex even after cortical firing was abolished optogenetically. We therefore propose that the oscillation is generated either within thalamic circuits, or by their interactions with their retinal inputs.

Our results are consistent with previous reports of narrowband oscillations in mouse visual cortex at frequencies close to 60 Hz ^25, 26^. Furthermore, the oscillation we study here might be homologous to an oscillation previously described in LGN of anesthetized cats ^23, 24, 43^, which showed extremely narrowband rhythmicity within single recordings, although the frequency of this oscillation varied between recordings, spanning a range of 45 to 114 Hz. In these cat recordings, the thalamic oscillation was highly synchronous with a retinal oscillation determined either by simultaneous recording, or by analysis of intracellular EPSPs in thalamus. If this proposed homology is correct, it would suggest that the narrowband gamma recorded here might reflect an oscillation generated in retina and transmitted through LGN to visual cortex. The fact that the power of this oscillation is increased by locomotion (as well as by stimulation of mesencephalic locomotor nuclei at levels too small to evoke locomotion ^26^) suggests that if the oscillation is indeed retinally-generated, its amplitude is likely to be modulated by thalamic circuitry.

What function might this narrowband gamma oscillation play in visual processing? Although a definitive answer to this question requires further work, the present data are sufficient to formulate two hypotheses.

A first hypothesis is that the narrow band gamma oscillation represents an “idling” rhythm. In this interpretation, narrowband gamma represents a default state of LGN activity when visual processing is not being carried out, similar to the proposed function of the visual alpha rhythm and the beta oscillation of the motor system. This hypothesis would explain why this oscillation is suppressed by increases in visual contrast.

In the second hypothesis, the narrow and broadband gamma oscillations would represent specific channels for thalamocortical and corticocortical communication. It has been proposed that oscillatory patterns with different frequency characteristics may allow routing of information between specific brain structures. For example, the coherence of slow gamma between hippocampal areas CA1 and CA3, and fast gamma between CA1 and entorhinal cortex ^13^, may indicate the existence of two separate transmission channels in the hippocampal formation. Analogously, the narrowband gamma oscillation may provide a specific channel for feedforward transmission from LGN to visual cortex, whereas broadband gamma – which is generated in multiple cortical regions - instead reflects the transmission of information within and between cortical circuits.

## Acknowledgements

ABS, MC and KDH are funded by the Wellcome Trust (grants 095668 and 095669) and by the Simons Collaboration in the Global Brain (grant 325512). KDH is also funded by the EPSRC (K015141). MRR and LB were supported by funds awarded to the Centre for Integrative Neuroscience (DFG EXC 307). ADL and KR performed the experiments in Massimo Scanziani’s lab with the support of the Howard Hughes Medical Institute and the Gatsby Charitable Foundation. KR was also supported by the National Science Foundation Graduate Research Program Fellowship. We thank R. Dulinskas for help with data collection for Figure S3. MC holds the GlaxoSmithKilne/Fight for Sight Chair in Visual Neuroscience.

## Supplementary Figures

**Figure. S1.**
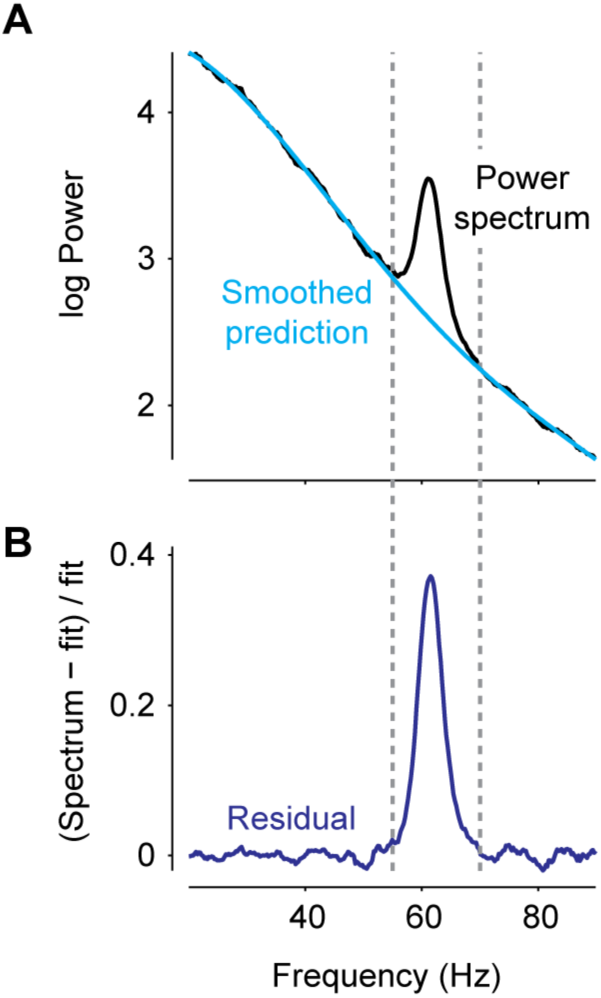
Supplementary Figure S1: Analysis of narrow-band oscillation. A. Example power spectrum from one recording session (Black). The power spectrum (excluding the narrow-band gamma region highlighted by the dotted lines) is fitted by a 4^th^ order polynomial (smoothed prediction, cyan). **B.** The residual spectrum is the difference between the actual data and the fitted function, divided by the fitted function. The peak frequency of narrow-band gamma was calculated as the position of the peak in the residual spectrum.

**Figure. S2.**
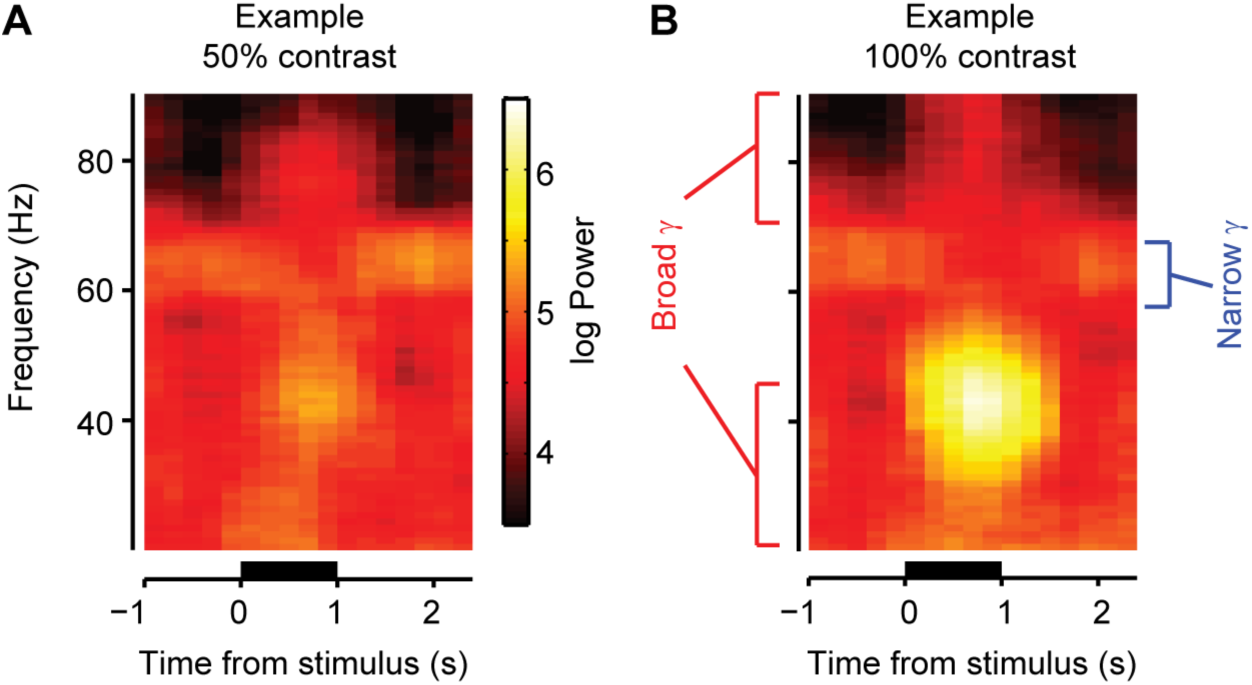
Supplementary Figure S2 (related to Figure 2): Broadband gamma rhythms are enhanced by visual contrast while the narrowband gamma oscillation is suppressed. A-B. Average LFP spectrogram triggered on presentation of a drifting grating stimulus at 50% contrast (**A**) and 100% contrast (**B**), in a single experiment. Bars below indicate the period of presentation of a 50^o^ sinusoidal drifting grating in the retinotopic location of the recorded area.

**Figure. S3.**
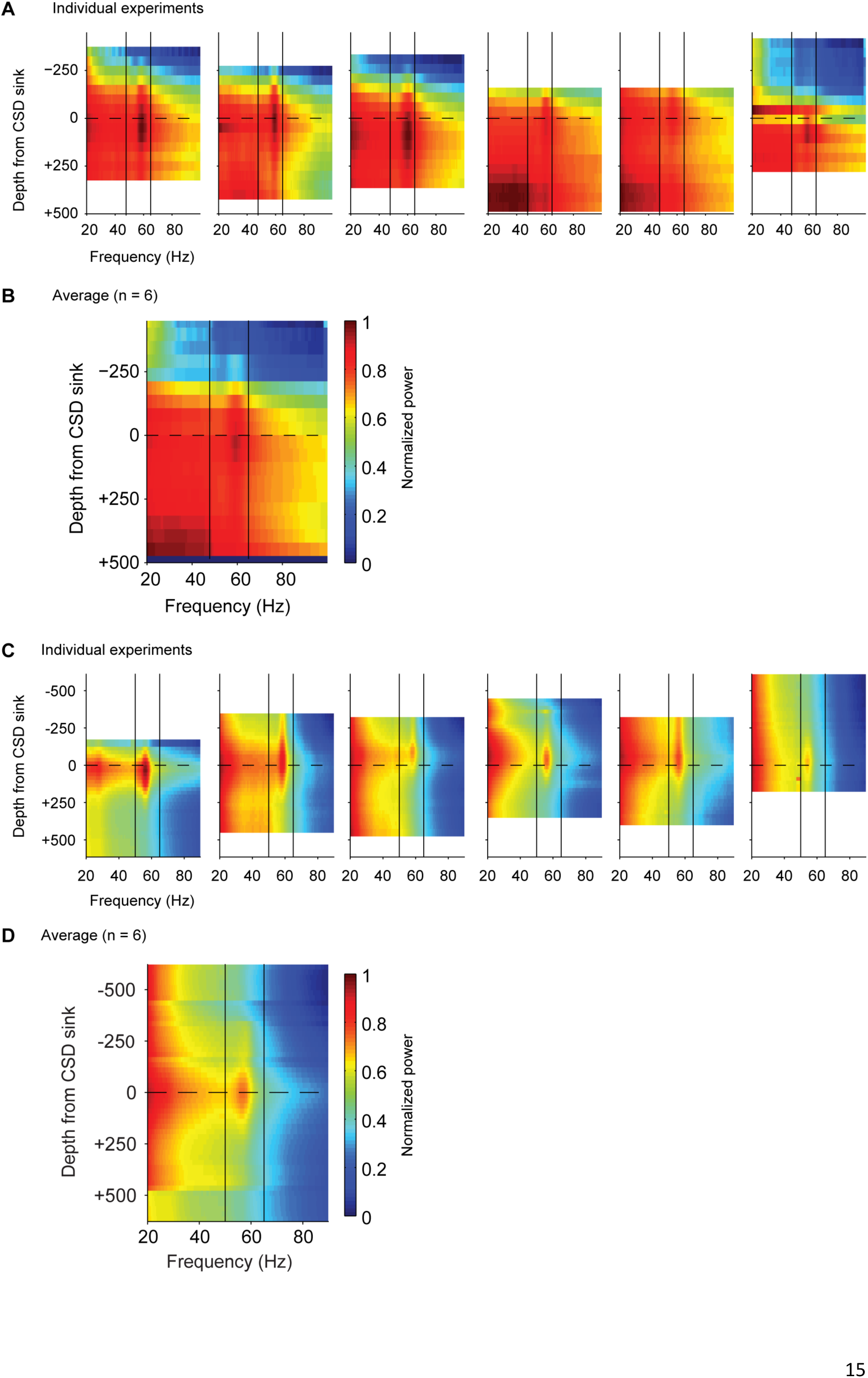
Supplementary Figure S3 (related to Figure 3): Narrowband gamma is highest in Layer 4 of V1 across experiments. A,C. The power across frequencies as a function of depth from the putative Layer 4 (peak of the CSD sink) across six experiments. The vertical lines mark the frequency range (50–65 Hz) of narrow-band gamma oscillation. **B,D.** The depth profile of narrow band gamma averaged across all experiments. Experiments in A-B were conducted using protocol A and experiments in C-D were conducted using protocol B. We thank R. Dulinskas for help with data collection (C, D).

**Figure. S4.**
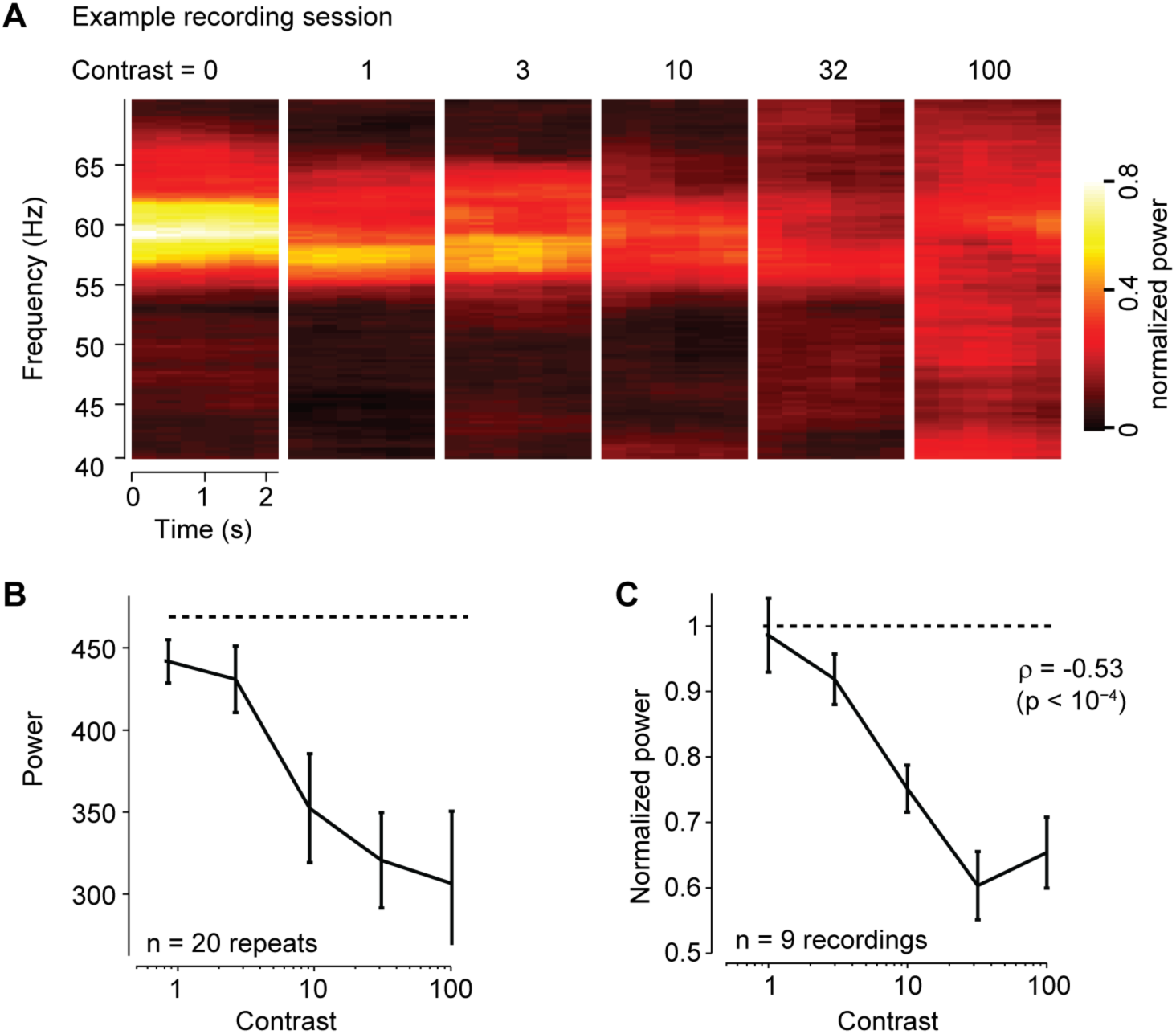
Supplementary Figure S4 (related to Figure 2 & Figure 5): Narrowband gamma oscillation in LGN spiking is suppressed by contrast. A. Average spectrogram of spiking activity of a single neuron, triggered on the start of full-screen grating stimuli (0%, 1%, 3%, 10%, 32% & 100% contrast, each contrast level repeated 20 times). The stimulus was presented for 2s, with a 3s inter-stimulus interval with 0% contrast (iso-luminant gray screen). **B.** Narrowband power decreases as a function of the contrast presented for the recording session shown in **A.** The dotted line shows the mean narrowband power at 0% contrast. **C.** Same as B, averaged across all the recording sessions (ρ = −0.53, p<10^−4^; n = 9 recordings). The peak narrowband power was normalized for each recording based on the power at 0% contrast.

**Figure. S5.**
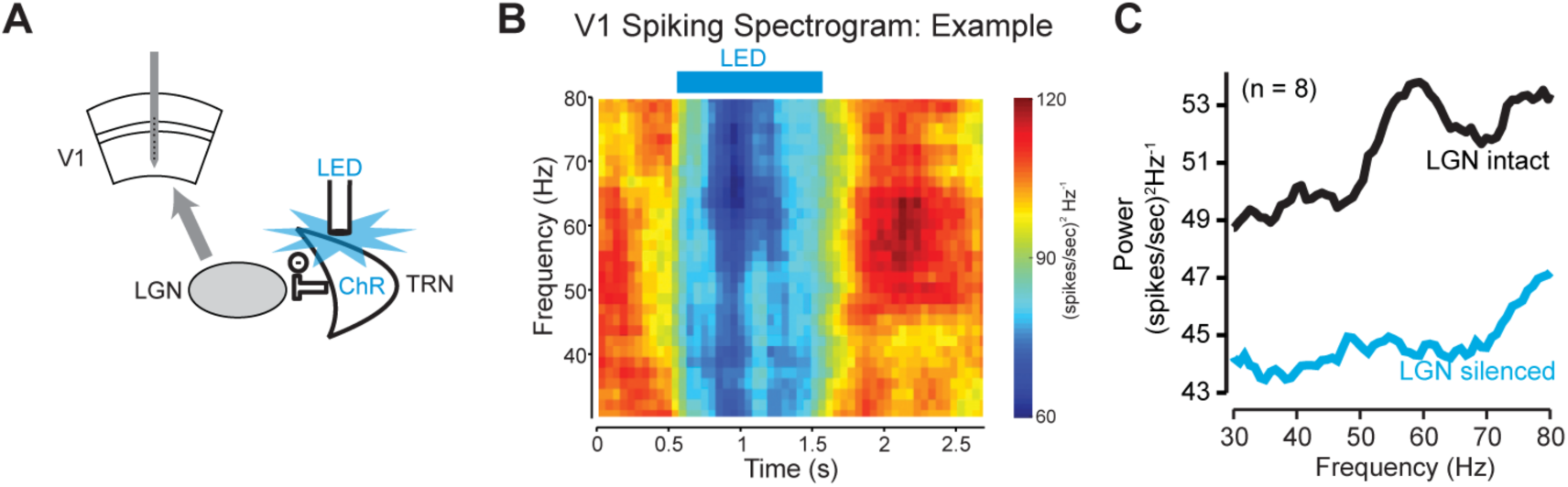
Supplementary Figure S5 (related to Figure 4): Narrowband gamma oscillation in V1 is suppressed by LGN inactivation. A. We recorded activity in V1 using a multi-electrode array (in awake mice), while inactivating the LGN by activating the TRN using an LED. **B.** Spiking spectrogram of V1 activity triggered on LED stimulation (Example experiment) **C.** The average power of V1 during (cyan) and after (black) LGN inactivation (n = 8 experiments). The narrowband gamma peak is abolished, together with a decrease in power at all frequencies.

**Figure. S6.**
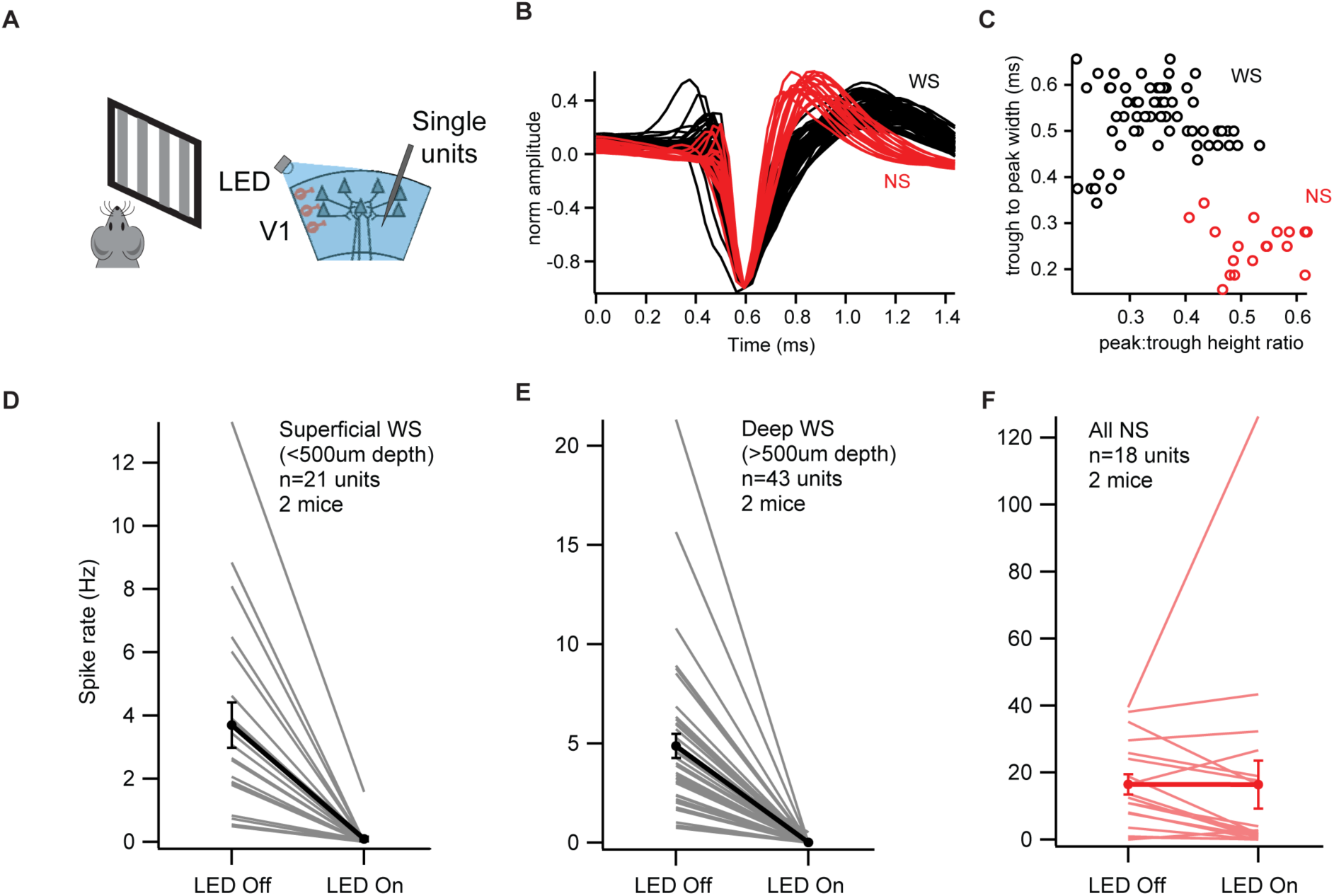
Supplementary Figure S6 (related to Figure 4): Cortical silencing is effective in suppressing excitatory activity in both superficial and deep layers of visual cortex. A. We recorded extracellular activity in V1 during the presentation of sinusoidal grating stimuli, with and without cortical silencing. **B.** Superimposed waveforms of the recorded neurons, color coded based on whether they were classified as wide spiking (WS, black) or narrow spiking (NS, red). **C.** The neurons were classified as WS or NS based on the peak:trough height ratio and trough to peak width (based on methods from Niell & Stryker, 2008). **D.** Turning on the LED suppressed the firing of all the WS neurons recorded from superficial layers (<500um; n = 21 neurons). E. Turning on the LED suppressed the firing of all the WS neurons recorded from deep layers (>500um; n = 43 neurons). **F.** The activity of NS neurons did not change significantly between the LED On and LED Off conditions (all depths; n = 18 neurons).

**Table 1.**
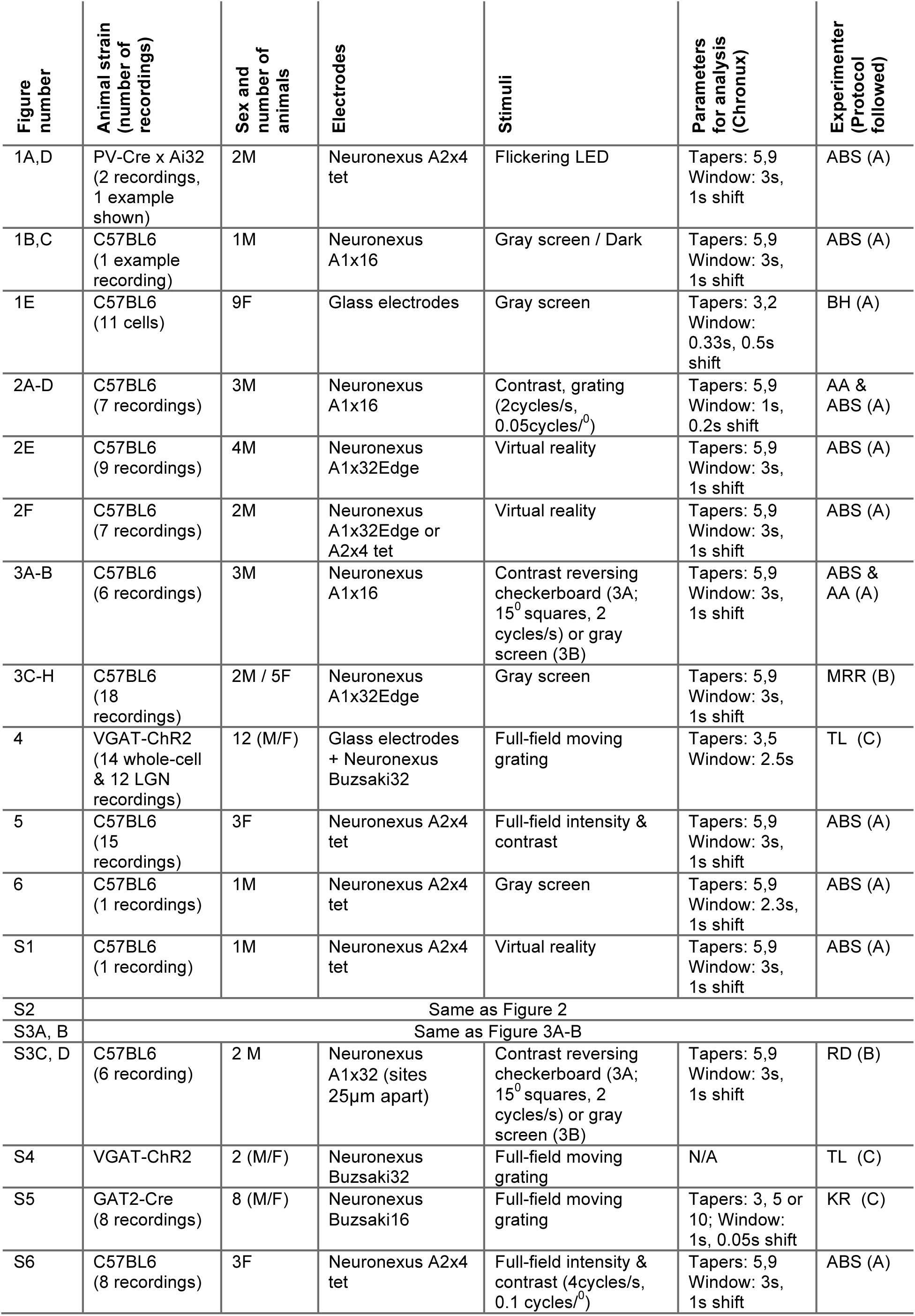
Supplementary Table 1: Details of recordings, stimuli and data analyses parameters

## Online methods

### Experimental methods

The experiments were performed in three different laboratories, following one of three protocols below. We list the protocol followed for each experiment in Supplementary Table 1.

<u>Protocol A</u> All procedures were conducted in accordance with the UK Animals Scientific Procedures Act (1986). Experiments were performed at University College London under personal and project licenses released by the Home Office following appropriate ethics review. We used both male and female mice of two strains: C57BL6 (17 animals) and PV-Ai32 (2 animals) (full listing in Supplementary Table 1). Experimental methods are described in detail in Ayaz et al. 2013, Haider et al. 2013 and Saleem et al. 2013^1-3^. Briefly, animals were chronically implanted with a custom-built head post and recording chamber (3–4 mm inner diameter) under isoflurane anesthesia. They were then allowed to recover for 3 days with oral analgesic Rimadyl. Following recovery, the animals were acclimatized for 3–4 days to the head fixation apparatus. A craniotomy was performed between 4 – 24 hrs prior to the first recording session. Whole-cell patch-clamp recordings of EPSCs and IPSCs were performed with a cesium-based internal solution, with QX-314 (0.5mM) and tetraethylammonium (TEA) (5mM) included to block voltage-gated Na+ and K+ conductances. Extracellular spikes were isolated using the KlustaView Suite^4^. Visual stimuli were presented on LCD monitors that were gamma corrected. For most experiments, the mean light intensity level presented on the monitors was ~50 cd/m^2^. The experiments in the dark were conducted making sure that all non-essential equipment was turned off, and essential equipment producing any light were covered with a filter (787 Marius Red; LEE filters) that only allows light that is not visible to mice. The light intensity level under these conditions was <10^−2^ cd/m^2^, below the sensitivity of the light meter. For experiments in Figure 5 and Supplementary Figure S6, we placed Fresnel lenses in front of the three monitors to ensure that the light intensity entering the eye is independent of the viewing angle on the LCD monitors. With Fresnel lenses the light intensity level could vary from ̴0.2 cd/m^2^ to ̴95 cd/m^2^.

<u>Protocol B</u> All procedures were performed on awake, adult mice and complied with the European Communities Council Directive 2010/63/EC and the German Law for Protection of Animals. All procedures were approved by the local authorities following appropriate ethics review. We used both male and female C57BL6 mice (full listing in Supplementary Table 1). Experimental methods are described in detail in Erisken et al, 2015^5^. Briefly, animals were chronically implanted with a custom-built head post and recording chamber (3–4 mm inner diameter) under isoflurane anesthesia. Animals were then allowed to recover for a minimum of 3 days during which they received analgesics (Carprofen, 5 mg/kg, s.c.) and antibiotics (Baytril, 5 mg/kg, s.c.). Following complete recovery, the animals were handled and acclimatized for a minimum of ̴1 week to head fixation and the spherical treadmill. One day before the first recording session, a craniotomy was performed under isoflurane anesthesia. Spikes were isolated using the Klusters suite ^6^ in combination with KlustaKwik ^7^. Visual stimuli were presented on LCD monitors that were gamma corrected. For most experiments, the mean light intensity level presented on the monitors was ̴50 Cd/m^2^.

<u>Protocol C</u> All procedures were conducted in accordance with the National Institutes of Health guidelines and with the approval of the Committee on Animal Care at UCSD (protocol S02160M). Animals were housed on a reverse light cycle in cages of four mice or less. At the time of electrophysiology, all animals were older than 3.5 weeks. Both male and female animals were used in an approximately equal ratio.

Experimental methods are described in detail in Lien et al. 2013 and Reinhold et al. 2015 ^8^, ^9^. Briefly, simultaneous V1 L4 intracellular whole-cell and LGN extracellular (Buzsaki32 probe, Neuronexus) recordings were performed in VGAT-ChR2 mice (JAX 014548 Zhao, et al) under urethane (1.5g/kg, IP)/chlorprothixene (2–4 mg/kg, IP) anesthesia. V1 neurons were held in voltage clamp at −70 mV to record excitatory currents. Full field drifting bar gratings (100% contrast, 2 Hz temporal frequency, 0.04 cycles/deg spatial frequency) were presented on a monitor in the right visual field for 2.3 s preceded by 0.7 s of a gray screen of mean light intensity. Optogenetic suppression of V1 was achieved by constant illumination of V1 with blue light from a 1mm fiber-coupled LED positioned several millimeters over the crantiotomy (455 nm, 20mW at the fiber tip, Doric) on interleaved trials starting 645 ms prior to grating onset and extinguishing after grating offset. For experiments inactivating the LGN, V1 extracellular (Buzsaki16 probe, Neuronexus) recordings were performed in awake Gad2-Cre mice (Jackson Labs stock number: 010802 ^10^) expressing Cre-dependent ChR2 (AAV2/1.CAGGS.flex.ChR2.tdTomato.SV40, Addgene 18917 ^11^, from the University of Pennsylvania viral vector core) stereotactically confined to the thalamic reticular nucleus (TRN). Mice were head-fixed but free to spontaneously run or rest on a circular treadmill. Full field drifting bar gratings (100% contrast, 2 Hz temporal frequency, 0.04 cycles/deg spatial frequency) were presented on a monitor in the right visual field for 3 s preceded by 2.5 s of a gray screen of mean light intensity. Optogenetic suppression of LGN was achieved by constant illumination of TRN with blue light from a 473 nm laser coupled to an optical fiber of diameter 200 microns, the tip of which was positioned just above the TRN (stereotactic coordinates of tip: [1,540 μm posterior, 2,235 μm lateral, 3,158 μm ventral to bregma], 10mW at the fiber tip). Laser illumination of the TRN on interleaved trials starting 0.2 ms after grating onset and lasting 1 second led to a 70% suppression ^9^ of spiking in LGN over the duration of illumination. Single units from both LGN and V1 recordings were isolated using UltraMegaSort ^12^. Visual stimuli were presented on an LCD monitor that was gamma corrected. The mean light intensity level presented on the monitors was 75 Cd/m^2^ for experiments in Figure 4.

## Analyses

### Spectral Analysis

We used the Chronux toolbox (http://chronux.org/), which is based on multi-taper methods, for spectral analyses. We measured the spectrogram of the local field potential using the function *mtspecgramc.* The power spectrum was calculated as the mean spectrogram across time. For spiking spectra, we used *mtcohgrampt* for the spike times (measured at 1 msec temporal resolution). We list the parameters we used for the spectral analysis (number of tapers and temporal window) in Supplementary Table 1. To calculate coherence across cells pairs, we used the Chronux function *mtcohgrampt.* To measure phase differences for any cell, we calculated its coherence with the pooled spike times from all other units. We calculated phase as the angle of the mean joint power across time. For Figure 5 and Supplementary Figure S6, we pooled the spike times across all units detected on a tetrode to calculate the spiking spectrogram. There were a total of 12 recording sessions in three animals where we recorded responses to varying levels of light intensity, two of these sessions had more than one tetrode in the LGN. There were a total of 8 recording sessions where we recorded responses to varying levels of contrast, one of which had more than one tetrode in the LGN.

### Residual power and peak frequency

We characterised the narrowband frequency using the residual power spectrum: the fractional increase in the power of the spectrum compared to a smoothed prediction (Supplementary Figure S1). We first calculated the smoothed prediction by fitting a 4^th^ order polynomial to the spectrum in the range 20–55 Hz and 70–90 Hz. This prediction generally captured the 1/f fall-off and the broadband gamma peak. We next calculated the residual spectrum at each frequency as:

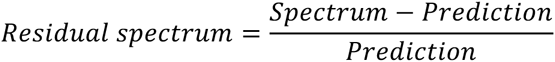

We define the narrowband frequency of any recording as the frequency with the maximum power of the residual spectrum in the range 55–70Hz. For spiking spectra from the LGN, we refer to the peak value of residual spectrum as the ‘residual gamma power’.

To statistically compare peak narrowband gamma frequencies between different LED flicker rates (Fig. 1D), we divided the recordings for each flicker rate into 5 equal subsets, computed a peak frequency in each subset, and compared these using paired t-tests for each pair of flicker rates.

### Co-modulation

To access how the power of different frequency components of the local field potentials varied together, we calculated the co-modulation: the correlation of the spectral power of the two frequencies across time. We measured this at all pairs of frequencies in the range of 20–90 Hz.

### Cross-correlation

When computing cross-correlograms for neurons from the same recording tetrode (Fig. 3F), the central bin was discarded and interpolated to avoid spike collision artifacts.

